## Supplementary Figures for "Protein absorption in the zebrafish gut is regulated by interactions between lysosome rich enterocytes and the microbiome"

### Supplemental Information

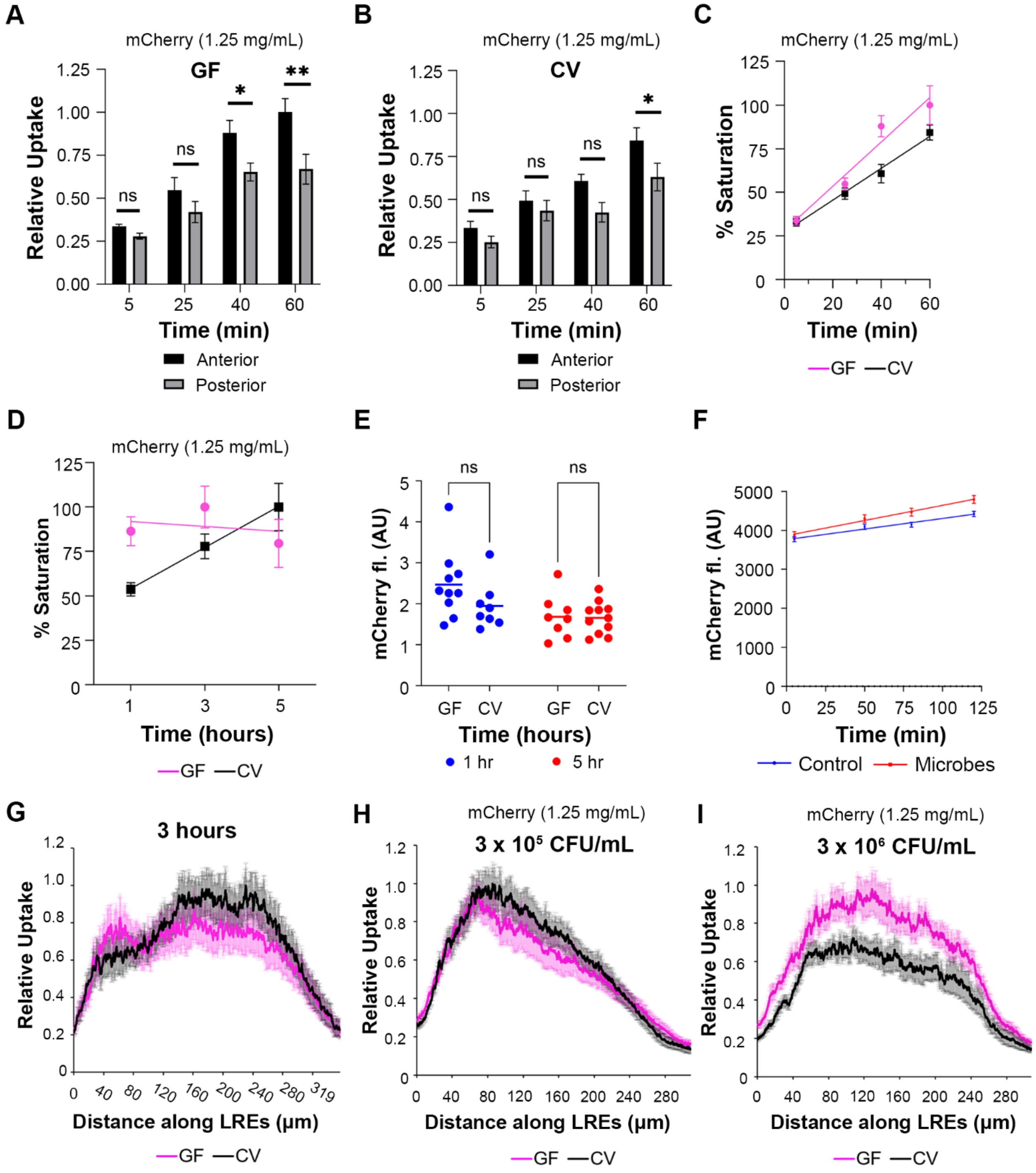


**Supplemental Figure 1. Regional differences and impacts of microbial density on cargo uptake by LREs.**

(A, B) Plots comparing normalized cargo uptake by anterior (50 - 150 μm) and posterior (150 - 250 μm) LRE regions in 6 dpf GF (A) and CV (B) larvae. The anterior LREs took up significantly more mCherry than posterior LREs at 40 minutes (2-way ANOVA, padj = 0.047, n = 9 - 11) and 60 minutes PG in GF larvae (2-way ANOVA, padj = 0.0013, n = 11). The anterior LREs took up significantly more mCherry than posterior LREs by 60 minutes post gavage in CV larvae (2-way ANOVA, padj = 0.043, n = 9 - 10).

(C) Plot showing % of maximal uptake (saturation) over time from 5 – 60 minutes post gavage. The anterior LREs of GF larvae (50 – 150 μm) took up mCherry at a significantly faster rate than CV larvae from 5 - 60 minutes post gavage (Simple linear regression, $p < 0.0001$, n = 9 - 11).
(D) Plot showing % of maximal uptake (saturation) over time. Between 1 – 5 hours post gavage, mCherry saturation remained constant in the anterior LREs (0 – 100 μm) GF larvae (Simple linear regression, p = 0.74, n = 8 - 10) but increased in CV larvae (Simple linear regression, p = 0.026, n = 8 - 11). The rate of mCherry accumulation was significantly different between GF and CV larvae (Simple linear regression, p = 0.0187, n = 8 - 11).
(E) Plot showing luminal mCherry fluorescence at 1 and 5 hours post gavage in GF and CV larvae. There was no difference in luminal fluorescence between conditions (2-way ANOVA, padj = 0.17, n = 8 – 11).
(F) Plot showing mCherry fluorescence (AU) over time in media containing zebrafish larva microbes or vehicle control. The change in mCherry fluorescence over time was not significantly different between zebrafish microbe and control media (Simple linear regression, p = 0.103, n = 8 - 10), showing the zebrafish microbiome did not degrade mCherry.
(G) Plot showing LY uptake in GF and CV larvae at 3 hours post gavage. There was no difference in LY fluorescence between GF and CV larvae (2-way ANOVA, p = 0.98, n = 9 - 11).
(H, I) Plots of normalized mCherry fluorescence intensity along the LRE region over time in 6 dpf GF and CV larvae. The LREs in GF and CV larvae took up the same amount of mCherry by 1 hour post gavage when the microbial density was $3 \times 10^5$ CFU/mL (2-way ANOVA, p = 0.18, n = 10) (H). GF larvae took up significantly more mCherry than CV larvae when the microbial density was $3 \times 10^6$ CFU/mL (2-way ANOVA, $p < 0.0001$, n = 10) (I).

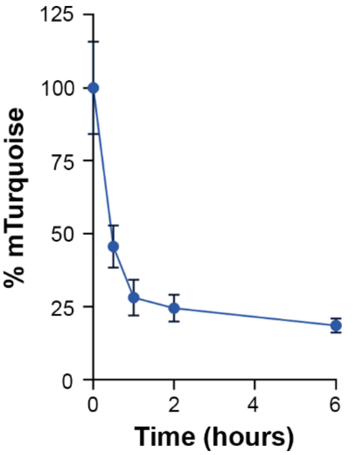


**Supplemental Figure 2. mTurquoise degradation kinetics in LREs.**
Plot showing mTurquoise degradation kinetics in LREs from 6 dpf, CV larvae over time. Over 75% was degraded within 2 hours.

**A**

| Cluster | GF | CV |
|---|---|---|
| Pharynx | 204 | 86 |
| Anterior enterocytes | 2,249 | 1,495 |
| Ileocytes | 229 | 264 |
| Anterior LREs | 131 | 199 |
| Posterior LREs | 359 | 381 |
| Cloaca 3 | 0 | 91 |
| Goblet | 267 | 226 |
| Enteroendocrine | 140 | 79 |
| Neuronal | 124 | 137 |
| Acinar | 70 | 33 |
| Best4/Otop2 enterocytes | 211 | 141 |
| Pharynx-cloaca 1 | 471 | 354 |
| Pharynx-cloaca 2 | 110 | 38 |
| Epidermis | 107 | 74 |
| Tuft and immune | 358 | 225 |
| Cluster 5 | 337 | 256 |
| Cluster 15 | 61 | 62 |

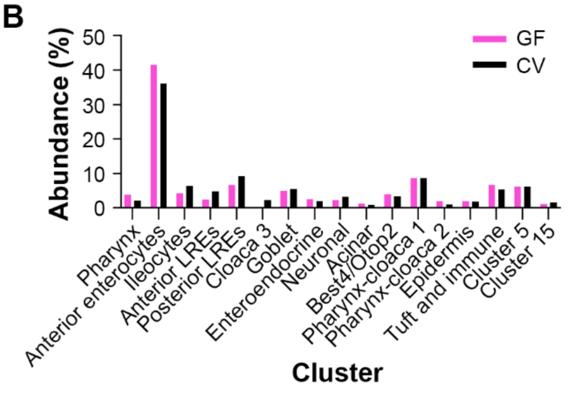


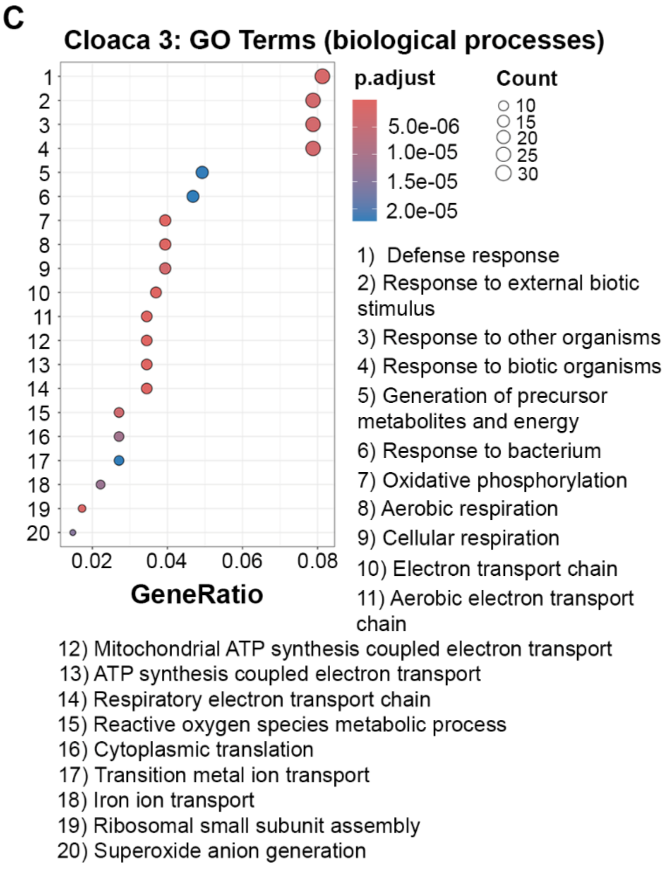


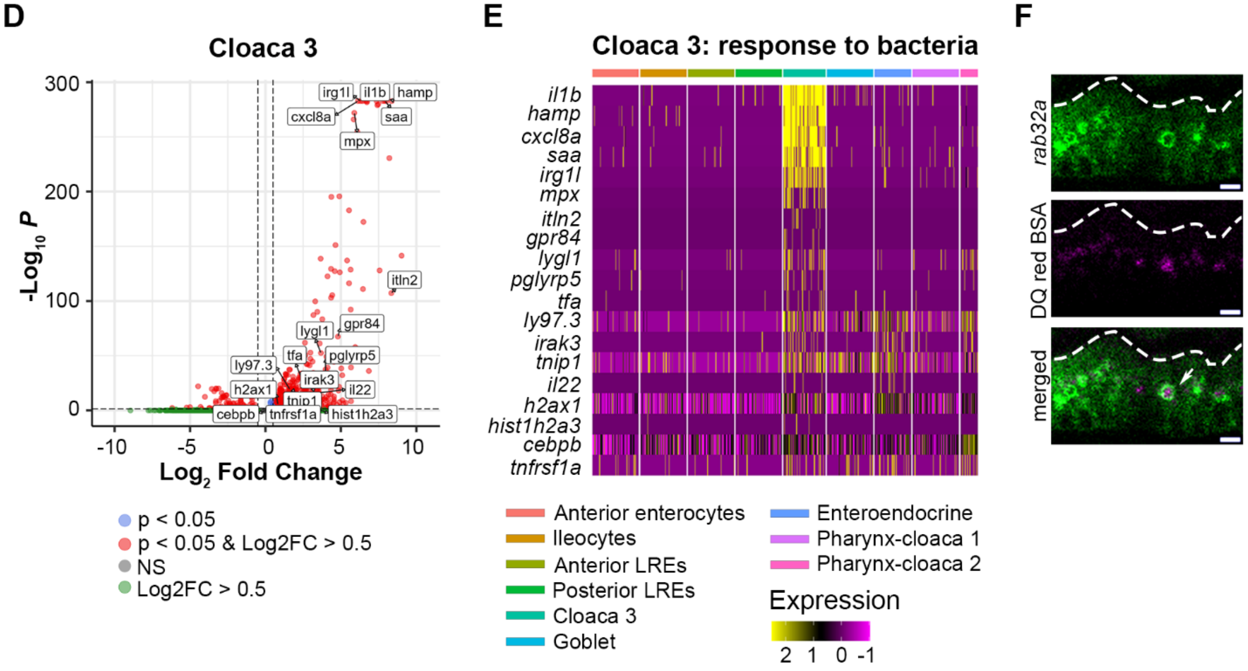


**Supplemental Figure 3. Cell counts and features of CV-specific Cloaca 3 cells.**
(A) Table showing the number of GF and CV cells per cluster in the scRNA-seq dataset.

(B) Bar plot showing the % abundance of cells that fell within each cluster in the GF and CV datasets. Both conditions showed comparable proportions of cells per cluster except for Cloaca 3, which only appeared in the CV dataset.
(C) Dot plot displays the GO terms that were upregulated in Cloaca 3 cells. The dot color indicates the adjusted p value for each GO term, while the dot size signifies the number of genes expressed in Cloaca 3 cells that fall into each GO term category. GeneRatio describes the proportion of genes associated with each GO term.
(D) Volcano plot illustrates markers upregulated in the Cloaca 3 cluster. Red points with log2FC > 0.5 are genes significantly upregulated in Cloaca 3 cells compared to other IECs. Labeled genes were categorized in the GO term "response to bacterium."
(E) Heatmap displays genes that were included in the "response to bacterium" GO term, which was upregulated in Cloaca 3. Cluster identity is indicated by the colored bars at the top of the heatmap. Expression level is shown by a color gradient, with yellow indicating the highest expression level.
(F) Live confocal microscopy images of LREs in 6 dpf larva expressing *GFP-rab32a* following gavage with DQ red BSA. DQ red BSA fluorescence was localized to the lysosome (arrow). Scale bars = 5 μm.

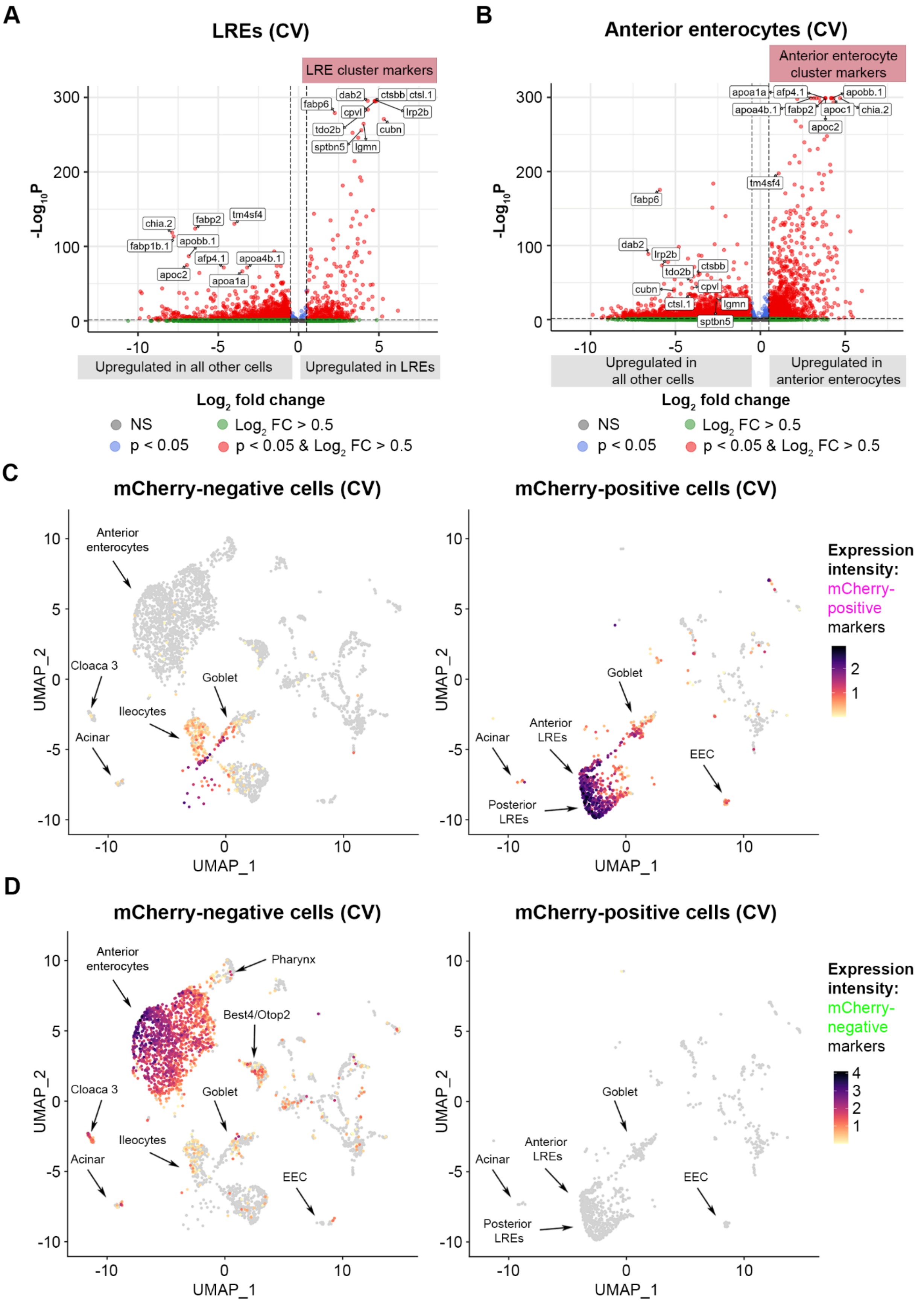

A
LREs (CV)
LRE cluster markers
-Log10 P
Upregulated in all other cells
Upregulated in LREs
Log2 fold change
NS
Log2 FC > 0.5
p < 0.05
p < 0.05 & Log2 FC > 0.5
B
Anterior enterocytes (CV)
Anterior enterocyte cluster markers
Upregulated in all other cells
Upregulated in anterior enterocytes
C
mCherry-negative cells (CV)
mCherry-positive cells (CV)
Anterior enterocytes
Cloaca 3
Ileocytes
Goblet
Acinar
Anterior LREs
Posterior LREs
EEC
UMAP_1
UMAP_2
Expression intensity: mCherry-positive markers
D
Pharynx
Best4/Otop2
Expression intensity: mCherry-negative markers

**Supplemental Figure 4. LRE and anterior enterocyte marker expression delineates mCherry-positive and mCherry-negative cells.**

(A) Volcano plot displays LRE marker genes in all cells within the CV dataset. The top differentially expressed genes (DEGs) between LREs and other cells are tagged. Genes with a red dot and log2FC > 0.5 are significant LRE cluster markers.

(B) Volcano plot displays anterior enterocyte marker genes in all cells within the CV dataset. The top DEGs between anterior enterocytes and other cells are tagged. Genes with a red dot and log2FC > 0.5 are anterior enterocyte cluster markers.

(C) UMAP projections show expression of the top DEGs in mCherry-positive cells compared to mCherry-negative cells (*ctsbb*, *lrp2b*, *ctsl.1*, *dab2*, *cpvl*, *cubn*, *tdo2b*, *lgmn*, *fabp6*, *sptbn5*) in the CV dataset. The cell color gradient intensity indicates the cumulative expression level of these genes. Left: UMAP projection displays all sorted mCherry-negative cells in the CV dataset. Right: UMAP projection displays sorted mCherry-positive cells in the CV dataset.

(D) UMAP projections show expression of the top DEGs in mCherry-negative cells compared to mCherry-positive cells (*fabp2*, *chia.2*, *fabp1b.1*, *apobb.1*, *apoa4b.1*, *apoa1a*, *tm4sf4*, *afp4.1*, *apoc2*, *apoc1*) within the CV dataset. The cell color gradient intensity indicates the cumulative expression level of these genes. Left: UMAP projection displays all sorted mCherry-negative cells in the CV dataset. Right: UMAP projection displays all sorted mCherry-positive cells in the CV dataset.

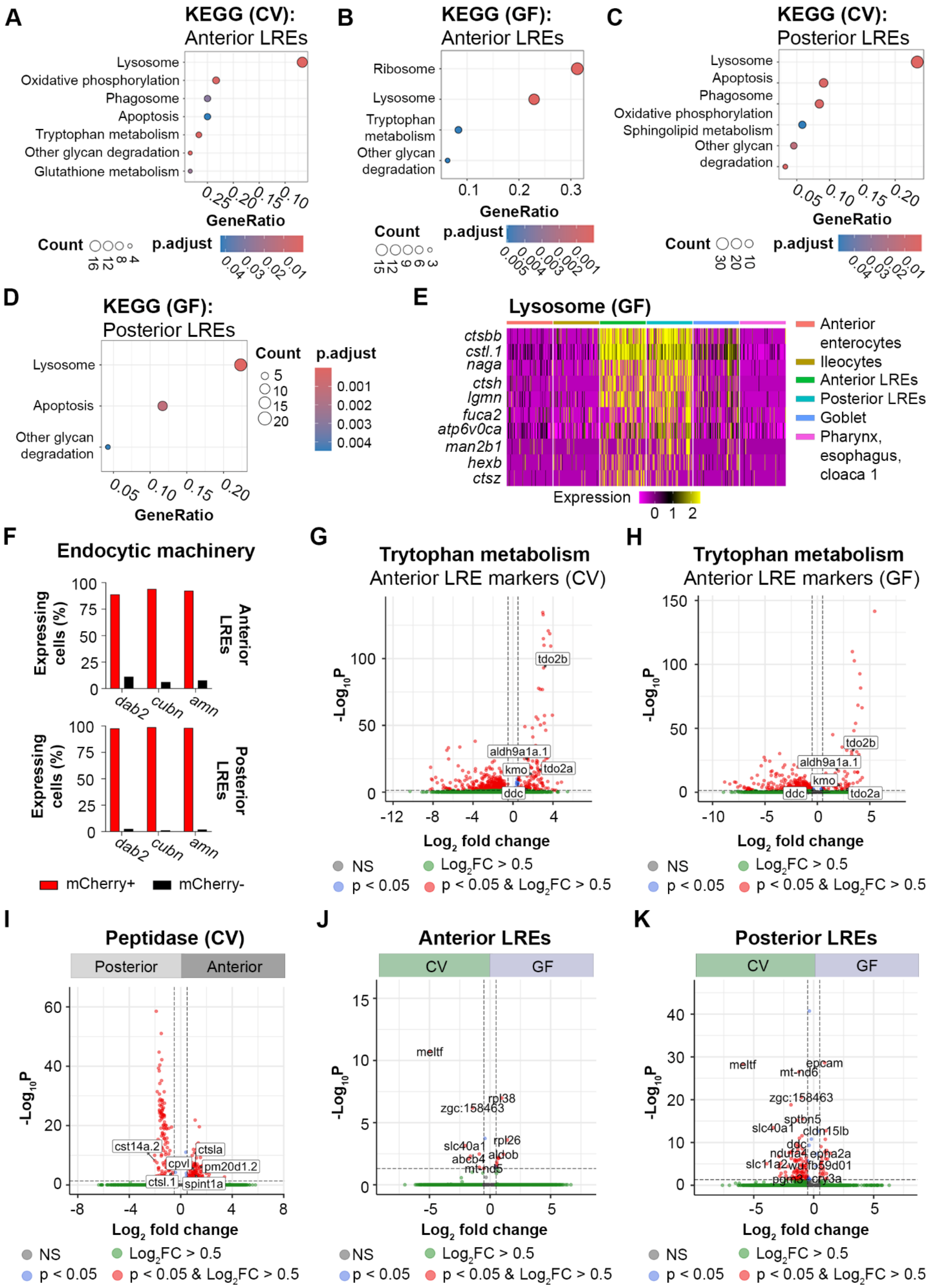
A
KEGG (CV): Anterior LREs
Lysosome
Oxidative phosphorylation
Phagosome
Apoptosis
Tryptophan metabolism
Other glycan degradation
Glutathione metabolism
GeneRatio
Count
p.adjust
B
KEGG (GF): Anterior LREs
Ribosome
Lysosome
Tryptophan metabolism
Other glycan degradation
GeneRatio
Count
p.adjust
C
KEGG (CV): Posterior LREs
Lysosome
Apoptosis
Phagosome
Oxidative phosphorylation
Sphingolipid metabolism
Other glycan degradation
GeneRatio
Count
p.adjust
D
KEGG (GF): Posterior LREs
Lysosome
Apoptosis
Other glycan degradation
GeneRatio
Count
p.adjust
E
Lysosome (GF)
ctsbb
cstl.1
naga
ctsh
lgmn
fuca2
atp6v0ca
man2b1
hexb
ctsz
Expression
Anterior enterocytes
Ileocytes
Anterior LREs
Posterior LREs
Goblet
Pharynx, esophagus, cloaca 1
F
Endocytic machinery
Expressing cells (%)
Anterior LREs
Posterior LREs
dab2
cubn
amn
mCherry+
mCherry-
G
Trytophan metabolism
Anterior LRE markers (CV)
tdo2b
aldh9a1a.1
kmo
tdo2a
ddc
Log2 fold change
-Log10P
NS
Log2FC > 0.5
p < 0.05
p < 0.05 & Log2FC > 0.5
H
Trytophan metabolism
Anterior LRE markers (GF)
tdo2b
aldh9a1a.1
kmo
ddc
tdo2a
Log2 fold change
-Log10P
NS
Log2FC > 0.5
p < 0.05
p < 0.05 & Log2FC > 0.5
I
Peptidase (CV)
Posterior
Anterior
cst14a.2
ctsla
cpvl
pm20d1.2
ctsl.1
spint1a
Log2 fold change
-Log10P
NS
Log2FC > 0.5
p < 0.05
p < 0.05 & Log2FC > 0.5
J
Anterior LREs
CV
GF
meltf
rpl38
zgc:158463
slc40a1
rpl26
abcb4
aldob
mt-nd5
Log2 fold change
-Log10P
NS
Log2FC > 0.5
p < 0.05
p < 0.05 & Log2FC > 0.5
K
Posterior LREs
CV
GF
meltf
epcam
mt-nd6
zgc:158463
sptbn5
slc40a1
cldn15lb
ddc
ndufa4
epha2a
slc11a2
wu:fb59d01
pgm3
cry3a
Log2 fold change
-Log10P
NS
Log2FC > 0.5
p < 0.05
p < 0.05 & Log2FC > 0.5

**Supplemental Figure 5. LREs show regional expression patterns and responses to the gut microbiome.**
(A) Dot plot shows KEGG pathways that were significantly upregulated in anterior LREs from the CV dataset. Dot color corresponds to the adjusted p value, while dot size describes the number of pathway genes expressed in anterior LREs.
(B) Dot plot of KEGG pathways that were significantly upregulated in anterior LREs of GF larvae.
(C) Dot plot of KEGG pathways that were significantly upregulated in the posterior LREs of CV larvae.
(D) Dot plot of KEGG pathways that were significantly upregulated in the posterior LREs of GF larvae.
(E) Heatmap showing expression of KEGG pathway lysosome genes by LREs, midgut cell types and Pharynx, esophagus, cloaca 1 cells in the GF larvae.
(F) Bar plots show the percentage of mCherry-positive and mCherry-negative cells that express components of the endocytic machinery in anterior LREs (top) and posterior LREs (bottom).
(G, H) Volcano plot shows differentially expressed genes between anterior LREs and all other cell clusters in the CV (G) and GF (H) datasets. Genes that play a role in tryptophan metabolism are labeled. Red points with log2FC > 0.5 are upregulated genes in anterior LREs from the CV (F) and GF (G) datasets.
(I) Volcano plot shows differentially expressed genes between anterior and posterior LREs in the CV dataset. Tagged genes are peptidases. Red points with log2FC > 0.5 and genes that were upregulated in anterior LREs, while red points with log2FC < -0.5 were upregulated genes in posterior LREs.
(J, K) Volcano plots show differentially expressed genes between GF and CV anterior (J) and posterior LREs (K). Genes marked with red points with log2FC > 0.5 were upregulated in the GF condition, while log2FC < 0.5 were upregulated in the CV condition.

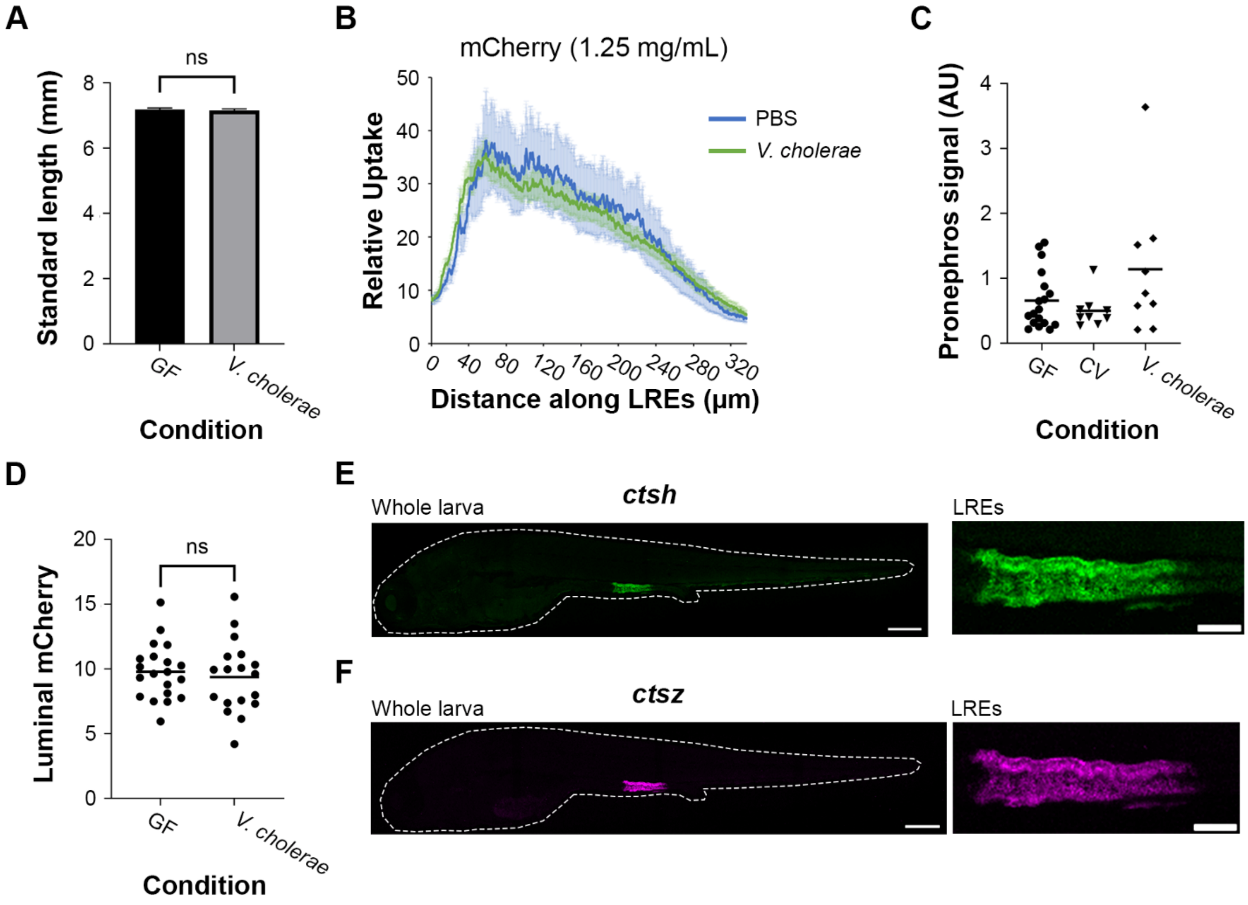


**Supplemental Figure 6. Long-term exposure to *V. cholerae* required to reduce protein uptake activity in LREs without affecting larval growth or protein availability.**
(A) Plot of standard length in larvae that were GF or monoassociated with *V. cholerae*. There was not a significant difference in standard length (Two-tailed T-test, p = 0.66, n = 18 – 22).

(B) Plot of mCherry uptake in larvae gavaged with PBS or live *V. cholerae*. There was no difference in the average mCherry uptake between the conditions (2-way ANOVA, $p = 0.527$, $n = 7 – 16$).
(C) Plot of mCherry fluorescence signal in pronephros of GF, CV and *V. cholerae*-colonized larvae. There was no significant difference between GF, CV and *V. cholerae* (1-way ANOVA, $p = 0.154$, $n = 9$).
(D) Plot of average luminal mCherry fluorescence in GF and *V. cholerae*-colonized larvae at 6 dpf by 1 hour PG. Luminal mCherry fluorescence was not significantly different (Two-tailed t-test, $p = 0.60$, $n = 18 – 20$).
(E-F) Confocal images of *ctsh* (E) and *ctsz* (F) HCR probe localization in whole larva and the LRE region. Larva is outlined with a dashed line. Whole larva scale = 200 µm. LRE scale = 50 µm.

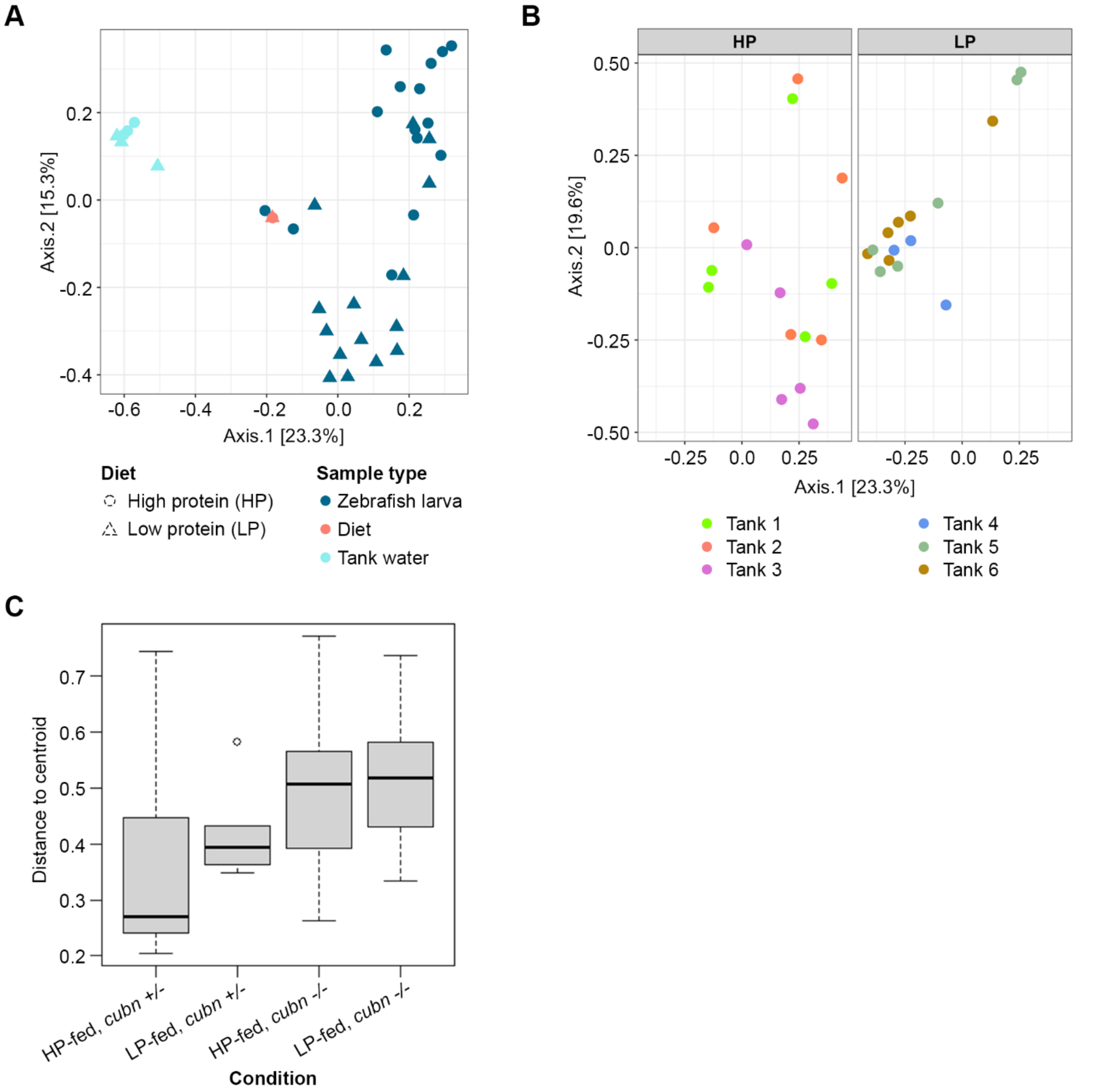


**Supplemental Figure 7. Microbiome assembly shaped by host environment.**
(A) MDS plot of Bray Curtis beta distance between sample types.
(B) MDS plot of Bray Curtis beta distance between zebrafish larvae fed the high protein (HP) or low protein (LP) diet. Point color indicates the tank that housed the zebrafish.
(C) Box and whisker plot of Bray Curtis beta dispersion between zebrafish larvae from each experimental condition.
